# When should coevolution among competitors promote coexistence versus exclusion?

**DOI:** 10.1101/2022.11.01.514799

**Authors:** Lucas A. Nell, Joseph S. Phillips, Anthony R. Ives

## Abstract

Coevolution of competitors can lead to niche partitioning promoting coexistence or to heightened conflicts promoting competitive exclusion. If both are possible, when should coevolution favor coexistence versus exclusion? We investigated this question with a general eco-evolutionary model in which species can reduce the interspecific competition they experience through evolutionary investments in two types of competitive traits: partitioning traits that promote coexistence and conflict traits that promote exclusion. We found that communities were generally mixed, consisting of species investing in both trait types or mixtures of species specializing in one type. For each species, its competitors’ abundances and investments determined its experienced competition, and stronger competition begot greater competitive trait investment. Species investing in conflict traits strengthened competition for other species both directly and indirectly, whereas partitioning traits only weakened competition via direct effects. Conflict traits were therefore the stronger driver of community-wide investments in all traits. However, species investing most in conflict traits experienced less competition, so they ultimately evolved least investment, making them most likely to be excluded by the next invader. Thus, coevolution may provide an open door for species that play nice and a revolving door of exclusion for those that do not.

## Introduction

Interactions among competing species can either promote or inhibit coexistence. Those that weaken interspecific competition relative to intraspecific competition should promote coexistence, while those that do the opposite should inhibit it (Chesson, 2000). Traits that promote coexistence often entail some form of niche partitioning, where species differ in the factors that affect their population growth rate (i.e., have different niches). Classic examples include species using different resources (Abrams and Rueffler, 2009; Macarthur and Levins,1967; Roughgarden, 1976). However, any trait affecting a population’s growth rate can serve as the basis for niche partitioning, including interactions with natural enemies (Abrams and Chen,2002; Ehrlich et al., 2017; Grover and Holt, 1998; Vandermeer and Maruca, 1998) and responses to environmental change (Armstrong and McGehee, 1976; Chesson, 1994; Chesson and Huntly,1997; Kremer and Klausmeier, 2017; Loreau, 1992; Pacala and Tilman, 1994). Alternatively, traits that increase species’ per capita effects on their competitors should inhibit coexistence. These typically occur as traits that cause conflict between a species and its competitors. For example, taller plants can more successfully access light and shade their shorter competitors, leading to an arms race among plants for increased height within the constraints imposed by the physiological cost of being tall (Falster and Westoby, 2003). Other examples of conflict traits include seed size (Fagerström and Westoby, 1997; Geritz et al., 1999), agonistic behavior (Brown,1971), and body size (Kisdi and Geritz, 2001). While investment in conflict traits does not necessarily lead to competitive exclusion (Falster and Westoby, 2003; Kisdi and Geritz, 2001), competitive exclusion can result when investment is sufficiently high.

Under what conditions should coevolution of competitive traits lead to coexistence versus exclusion? This depends on the extent to which species invest in partitioning traits (those that reduce per capita interspecific competition on both itself and its competitors) versus conflict traits (those that reduce per capita interspecific competition on itself while increasing competition on its competitors). Most theoretical studies focus on partitioning traits taking a specific form, such as traits affecting resource use (e.g., Roughgarden, 1976; Shoresh et al., 2008) or apparent competition (e.g., Abrams, 1998; Abrams and Chen, 2002; Schreiber et al., 2011). Although some studies have explored the effect of coevolution on competitors’ traits using more general forms of competitive interactions (e.g., Abrams et al., 2013; Pastore et al., 2021; Vasseur et al., 2011), these models consider only the evolution of partitioning traits. Yet, both partitioning and conflict traits are likely ubiquitous in competitive communities. In tropical forests, for example, trees have evolved to better capture light (and thereby shade competitors), but many mechanisms, such as species-specific responses to herbivores, lead to partitioning traits (Wright, 2002). Additionally, although coevolution of competitors is most often thought to promote niche partitioning (Pfennig and Pfennig, 2012), experimental evidence has shown that either niche partitioning (Schluter, 1994; Zuppinger-Dingley et al., 2014) or exclusion-promoting traits (Germain et al., 2020; Hart et al., 2019; Miller et al., 2014; Zhao et al., 2016) can evolve in response to competition. Therefore, understanding how coevolution might lead to coexistence or exclusion of competing species requires models that include both partitioning and conflict traits.

Here, we use a general model of coevolving competitors to evaluate how partitioning and conflict traits evolve, and how coevolution affects the structure of communities. In the model, each species can reduce the interspecific competition it experiences by investing in partitioning and/or conflict traits, and the level of investment evolves through selection. Investment in partitioning traits by species *i* reduces the competitive effect that it has on all other competitors. Investment in conflict traits by species i increases the competitive effect that species i has on all other competitors. Investment incurs a cost to a species’ per capita growth rate that does not directly involve the competitive effects. We take a quantitative genetics approach to model selection on trait investment, and the model allows both additive-genetic and non-heritable variation. For additive-genetic trait variation, the costs for simultaneous investments in both partitioning and conflict traits can be non-additive. In the sub-additive case, the overall cost of investing in both partitioning and conflict traits is less than the summed costs of investing in each trait type separately, while for the super-additive case the overall cost is greater.

In previous models that focus on coexistence (e.g., Pastore et al., 2021; Taper and Case,1992), species reduce interspecific competition by evolving traits that determine usage along a resource gradient. Evolution away from the optimal value incurs a cost, and per capita effects of one species on another are proportional to the difference in their resource-uptake traits. Our model differs in that investments are modeled only in terms of their effects on competition experienced by the species under selection and its competitors, which encompasses competitive interactions in a more-general way. The costs of investment are exacted through decreases in density-independent per capita population growth rates for both partitioning and conflict traits, making it possible to compare the evolution of both types of traits. Because the traits are not conceptually tied to a specific mechanism underlying competition, the specific traits may differ among species. For example, competition for light and apparent competition via natural enemies could both be simultaneously included in our model. Treating traits as abstractions allows general insights into how conflict and partitioning traits will coevolve. Furthermore, the coevolution of traits will determine the frequency and magnitudes of partitioning and conflicts among species, and hence the competitive structure of communities.

Our model includes both ecological (population abundances) and evolutionary (trait values) variables, and therefore it is an ecological-evolutionary model. For presenting results, we rely primarily on simulations, although these are paralleled by analytical solutions provided in Appendix A. We focus on two questions. What determines the magnitude of evolutionary investment in partitioning and conflict traits in a community? And what types of communities are the end product of coevolution, communities dominated by species showing partitioning traits, species showing conflict traits, or species showing both?

## Methods

### Ecological dynamics

We used a discrete-time, modified Lotka–Volterra competition model similar to that used by Northfield and Ives (2013). Communities consist of *n* species, and each species *i* can invest in both partitioning (*p_i_*) and conflict (*x_i_*) traits. A species’ per capita population growth rate is

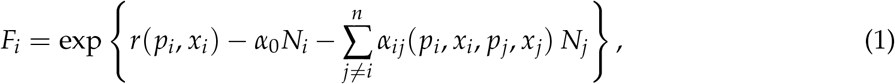

where *N_i_* is the population density of species *i*, and *α*_0_ is the strength of intraspecific competition which is fixed to the same value for all species and unchanged by trait evolution. The function *r*(*p_i_*, *x_i_*) is the per capita population growth rate of focal species i in the absence of any intra- or interspecific competition. This function describes the direct fitness costs of investment as a reduction from the baseline per capita population growth rate *r*_0_:

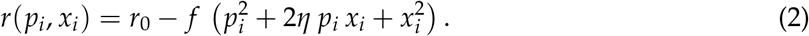

Here, *f* is a coefficient giving the cost of investment in a single trait type, and *η* governs the non-additivity of costs for investing in both suites simultaneously. When *η* > 0, costs are super-additive, and when *η* < 0, costs are sub-additive. Super-additive tradeoffs occur when investment in one suite of traits makes investment in the other suite of traits more costly (Garland et al., 2022). This might occur, for example, if competition for light selects for greater growth in plants, which then increases the cost of chemical defenses to specialist herbivores. Sub-additive costs could be caused by shared biochemical/physiological pathways that make investment in one type of trait also increase the effects of the other type (Garland and Carter,1994; Garland et al., 2022). Sub-additive costs could also arise when partitioning and conflict traits are correlated through an underlying trait (Strauss and Irwin, 2004), and therefore the evolution of one type of trait reduces the cost of the other. For example, an increase in body size could both improve the ability of a species in agonistic competition and allow it to feed on larger prey, thereby simultaneously leading to resource partitioning.

The term *α_ij_*(*p_i_*, *x_i_*, *p_j_*, *x_j_*) gives the pairwise interspecific competition coefficient, which is a function of the investments in partitioning and conflict traits for species *i* and *j*. We assume this has the form

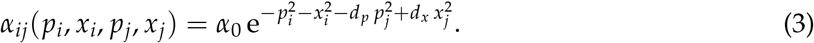

Interspecific competition is scaled by *α*_0_, such that the effects of investment in partitioning and conflict traits are defined relative to the fixed strength of intraspecific competition. The parameters *d_p_* and *d_x_* correspond to partitioning and conflict traits, respectively, and determine how strongly investments in each suite of traits in one species affect competition experienced by other species. With increasing values of *d_p_*, investment by species *i* in partitioning traits has a stronger negative effect on the interspecific competition experienced by other species. With increasing values of *d_x_*, investment by species *i* in conflict traits has a stronger positive effect on competition experienced by all other species. Values of *d_p_* and *d_x_* only affect a focal species *i* through the competition it experiences from other species *j* = *i*.

For both costs (eq. 2) and benefits (eq. 3), the functions giving the effects of trait investment on fitness are concave so that a single fitness peak exists for each suite of traits. We did this because our goal was not to model the diversification process, and because this made the equations more mathematically tractable, preventing runaway evolution to infinite trait values. Investments are constrained to be non-negative by passing the equations for *p*_*t*+1_ and *x*_*t*+1_ (eq. 4 below) through a ramp function. Details are given in Appendix A.

### Evolutionary dynamics

We used a quantitative genetics framework for modeling coevolution of traits by different species. We assumed that both *p_i_* and *x_i_* represent means for species *i* and that their among-individual distributions are symmetrical with additive genetic variance 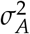. Assuming also that 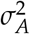 is relatively small (Abrams, 2001; Abrams et al., 1993; Iwasa et al., 1991), the evolution of traits are given by the equations

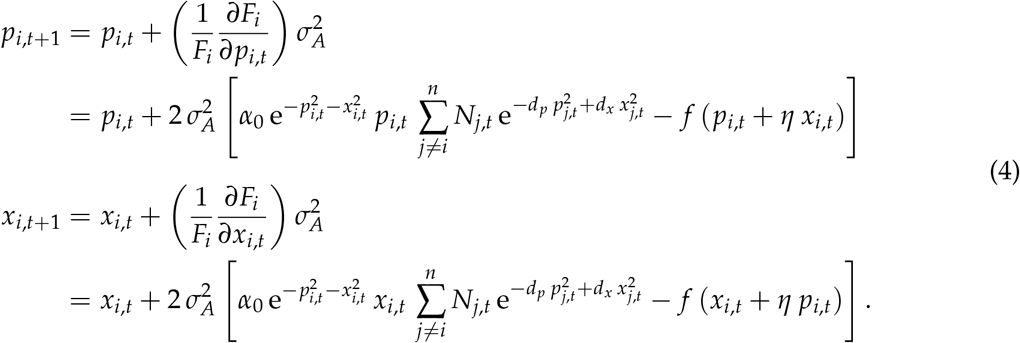

### Stability

The ecological and evolutionary stationary states produced by our model can be convergence stable and/or ESS stable (Geritz et al., 1998). A stationary (i.e., time-invariant) state is convergence stable if a nearby state evolves towards the stationary state. A stationary state is ESS-stable if, when occupied by the system, any mutant away from the state has lower fitness than those phenotypes at the stationary state. Convergence-stability does not imply ESS-stability, and vice versa (Geritz et al., 1998). Because we iterated the model until abundance and trait values reached a stationary state, the stationary state is convergence stable. To check for ESS convergence simultaneously for abundance and trait variables, we computed the Jacobian matrices for the abundances and investments of each species (eq. B5). The dominant eigenvalue of this matrix (*λ*) corresponds to a stationary state that is stable when ||*λ*|| < 1, neutrally stable when ||*λ*|| = 1, and unstable when ||*λ*|| > 1. Analytical expressions for the derivatives are shown in Appendix B.

### Non-heritable variation

In our model, changes in trait investments are driven by selection on additive genetic variance. However, in natural populations many factors other than additive genetic variance are likely to influence trait values that underpin competition, most notably plasticity (Miner et al., 2005). To explore the effects of non-heritable variation on trait coevolution, we modeled the expected values of the phenotypes 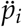 and *ẍ_i_* as the products of the latent additive-genetic components (*p_i_* and *x_i_*) and log-normal perturbations, such that 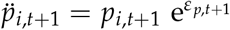 and *ẍ*_*i,t*+ 1_ = *x_*i, t*+1_*e^*ε*_*x,t*+1_^, where *ε*_*p,t*+1_ and *ε*_*x,t*+1_ are generated from normal distributions with means of 0 and variances 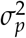 and 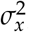, respectively. Because phenotypes affect both competition and associated costs, they are used to calculate all species’ fitness values and the effects of investments on competition. Genotypes change through time according to the effect of phenotypes on per-capita growth:

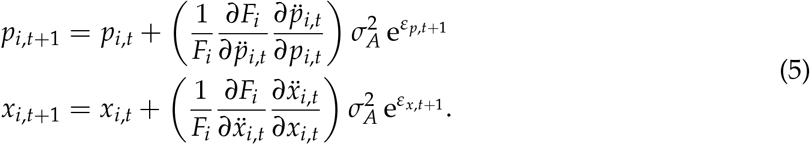

where 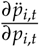 and 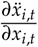 equal 1. We modeled non-heritable variation as simple perturbations because we were interested in the role of this variation per se—and not, for example, adaptive plasticity—and because non-heritable variation is often not adaptive (Ghalambor et al., 2007;Miner et al., 2005). The form of non-heritable variation we modeled could be produced by plasticity induced by stochastic environmental change. For example, temperature can change both nutritional demands and foraging behavior, so the costs and benefits of partitioning and conflict traits will likely depend on temperature.

### Simulations

We used simulated communities with 2 to 4 species to explore the behavior of the model. We do not present more species-rich communities because under the conditions we simulated, 2–4 species were sufficient to illustrate all of the qualitative outcomes of the model: for simulations with more species, species clustered into groups that were represented in our simulations of 2–4 species. To highlight different qualitative outcomes, we assumed symmetry among species by setting parameters *r*_0_, *α*_0_, and *f* to the same values for all species; using different values for different species changes the quantitative but not qualitative outcomes. All species started with an abundance of 1 and were considered extinct if their abundance fell below 1. We ran simulations long enough for communities to reach stationarity (≥ 10,000 time steps). Our R package `sauron` (https://anonymous.4open.science/r/sauron-F8CC) contains the code for the simulations that uses a combination of R (R Core Team, 2022) and C++, the latter via the Rcpp and RcppArmadillo packages (Eddelbuettel, 2013; Eddelbuettel and Sanderson, 2014; Sanderson and Curtin, 2016).

## Results

Our model of ecological and evolutionary dynamics investigates selection for partitioning and conflict traits, and how selection structures competition in the resulting community. To illustrate the central differences between partitioning and conflict traits, consider the case of two species and the effects on species B when species A changes its investment in either a partitioning or a conflict trait (fig. 1). To clarify this illustration, we fix the abundance and investment level of species A, rather than allow them to change as described above. When species A invests in a partitioning trait, the abundance of species B increases (fig. 1B), because greater investment in partitioning by species A reduces competition experienced by species B. With reduced competition, selection favors lower investment by species B. In contrast, for a conflict trait increased investment by species A causes a decrease in abundance of species B, and species B experiences selection for increased investment in the conflict trait (fig. 1C). Thus, investment in partitioning and conflict traits by species A have opposite effects on the abundance and selection for species B.

**Figure 1.**
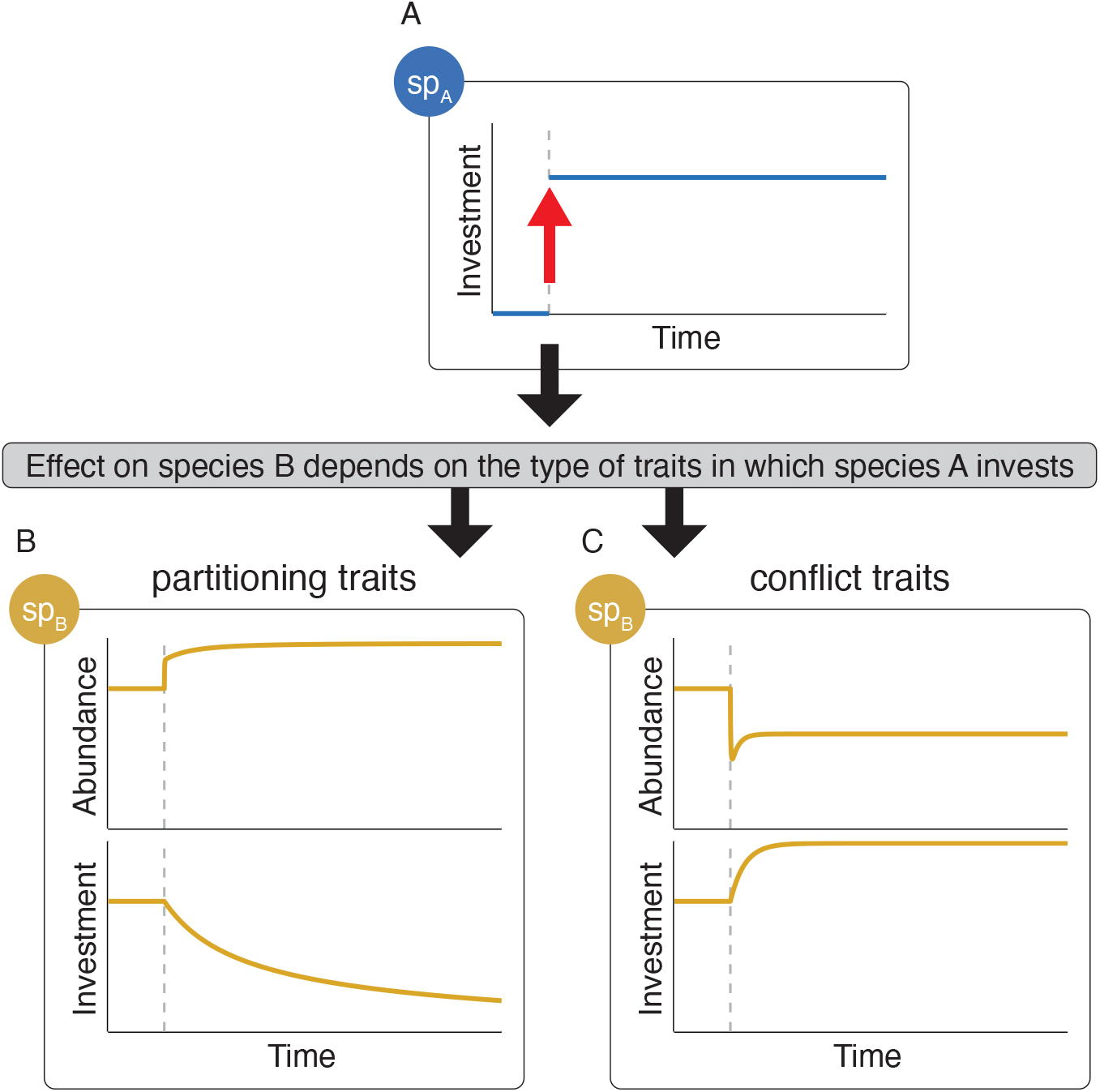
Simplified scenario demonstrating how investment by one species affects other species in the community. We start with a two-species community where the abundance and investments of species B are affected by species A and change though time as specified by the model. To show direct effects without feedbacks that complicate interpretation, these simulations have two differences from those throughout the rest of the paper: First, the abundance and investments of species A are constant and not affected by species B. Second, there is only one suite of traits. (A) If we perturb the system by instantly increasing the investment by species A, the effects this has on species B depend on the type of traits species A invests in. (B) If species A invests in partitioning traits, greater investment by A weakens the interspecific competition experienced by species B and causes a rapid initial increase in the abundance of B. Because reduced competition makes investment less beneficial, species B decreases its investment. The reduced costs of investing cause the slow increase in abundance of species B. (C) If species A invests in conflict traits, greater investment by A strengthens the interspecific competition experienced by species B and causes a rapid initial decrease in the abundance of B. Species B then evolves greater investment to reduce the effects of competition from A, which allows species B to increase in abundance after the initial decrease.

### Investment-cost additivity

We first show how different types of communities arise due to sub- and super-additive investment costs (fig. 2). We simulated 2-species communities where investment costs were either sub-additive (*η* =-0.6), additive (*η* =0), or super-additive (*η* =0.6). We added both species at the same time to avoid effects of evolution before interspecific competition starts. We set both suites of traits to be nearly neutral (*d_x_* = *d_p_* = 10^-6^) to reduce the effects of species’ investments on each other, which helps to highlight the effects of cost additivity only. The starting investments by all species were generated from a uniform distribution between 0 and 0.5. Sub-additive costs resulted in species evolving to a single point in investment space where each invests in both conflict and partitioning traits equally. Super-additive costs resulted in two alternative stable investment states: species evolved to invest in only partitioning or only conflict traits depending on their investments at the start of the simulations; different species could occupy different investment states even within the same community. Additive costs resulted in an intermediate case, with a neutrally stable ring given by 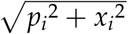 containing an infinite number of investment strategies with the same fitness; like the super-additive case, species within the same community can occupy different states. Because the total investment 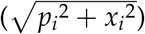 defining a ring is proportional to both the costs (eq. 2) and benefits (eq. 3) of investment, the ring observed in the additive case is also present in the other two cases. For sub-additive and super-additive costs, species trait values evolve towards the 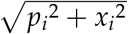 ring and then either converge to equal investment in partitioning and conflict traits (sub-additive) or diverge to investment in only partitioning or only conflict traits (super-additive) (fig. S1). Furthermore, larger magnitudes of *η* cause more rapid evolution around the ring (fig. S2). These results can be derived analytically (eqs. A3–A5).

**Figure 2.**
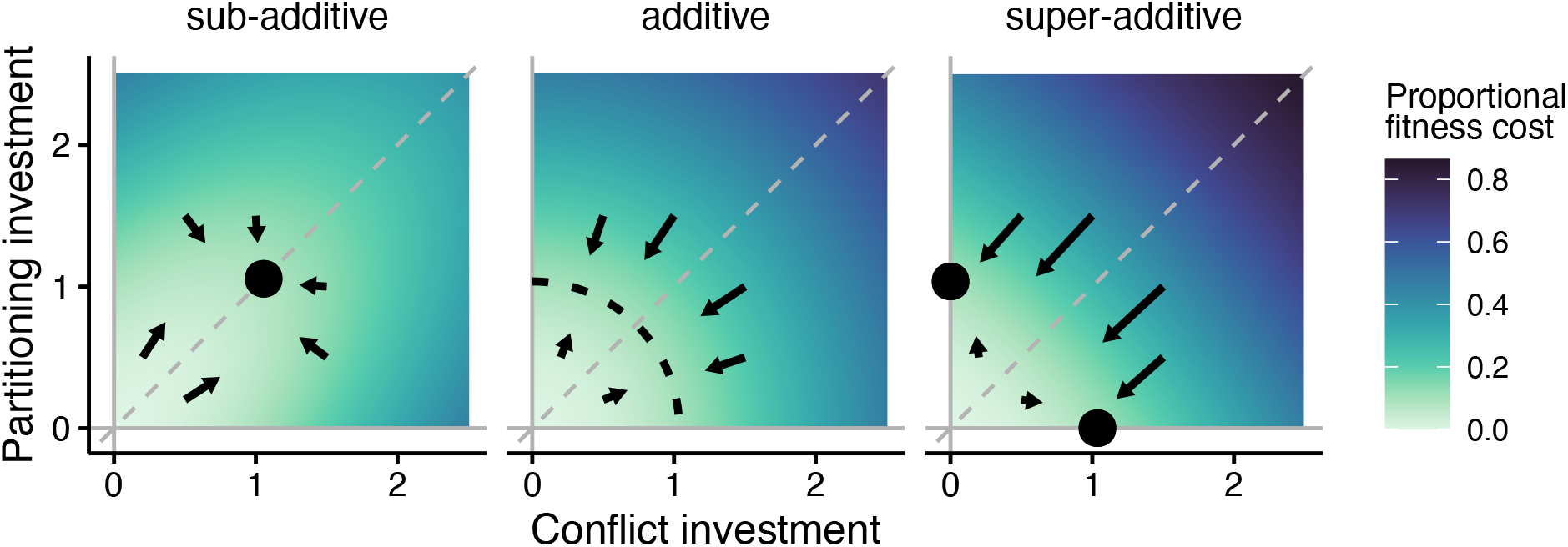
Costs of investing (background color) and unique investment states for 2-species communities (points and dotted line) for sub-additive (*η* < 0), additive (*η* = 0), or super-additive (*η* >0) investment costs. The dashed line in the additive case shows the location of the neutrally stable ring. Background color indicates the proportional reduction from the maximum per capita growth without competition 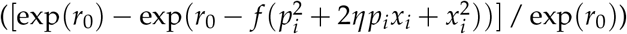, so darker colors indicate a greater cost of investment. Arrows show the direction and strength of selection (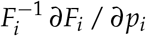 and 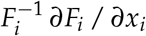) if a species at the nearest equilibrium point is perturbed to the arrow’s starting location.

Sub-additivity and super-additivity generate communities that differ in structure. At the community scale, sub-additive investment costs allow for only one possible stable configuration, where all species invest in both suites. For super-additive costs, there are three stable 2-species configurations: both species investing in partitioning traits, both investing in conflict traits, and one species investing in each. There are infinite community configurations for additive costs.

### Non-heritable variation

Because non-heritable variation can alter the effects of selection, we next explored the effects of non-heritable variation using simulations in which communities contained four species each with an initial total investment 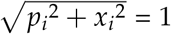. Two species had initial investments favoring partitioning traits, and the other two had higher initial investments in conflict traits (fig. 3). We considered three levels of non-heritable variation: non-existent (*σ_p_* = *σ_x_* = 0), low (*σ_p_* = *σ_x_* = 0.11), or high (*σ_p_* = *σ_x_* = 0.2). We crossed these levels to give nine permutations of non-heritable variation, for example, *σ_p_* = 0.2 and *σ_x_* = 0 in the top-left panels of figs. 3A, 3B, and 3C (where the non-heritable variation is given in the gray distributions on the sides of the panels). For the case when non-heritable variation affects partitioning and conflict traits the same (figs. 3A, 3B, 3C, subpanels along the diagonal), adding non-heritable variation causes the additive and super-additive cases to behave like the sub-additive case, whereby the neutrally stable ring and alternative stable states are replaced by a single stable state with investment in both partitioning and conflict traits. Furthermore, non-heritable variation reduces the efficacy of selection for a given trait; for example, when non-heritable variation for the partitioning trait is greater than for the conflict trait, the stable investment state has higher investment in the conflict trait. This is a manifestation of the classic result in quantitative genetics whereby the efficacy of selection is scaled by the additive-genetic (i.e., heritable) variance relative to the non-heritable variance (Barton, 2017; Fisher, 1930; Wright, 1931). These results are derived analytically in Appendix A (eq. A6). However, our results for non-heritable variation only apply for weakly super-additive traits. When costs are strongly non-additive (*η* = −0.6 or *η* = 0.6), non-heritable variation has little effect (fig. S3).

**Figure 3.**
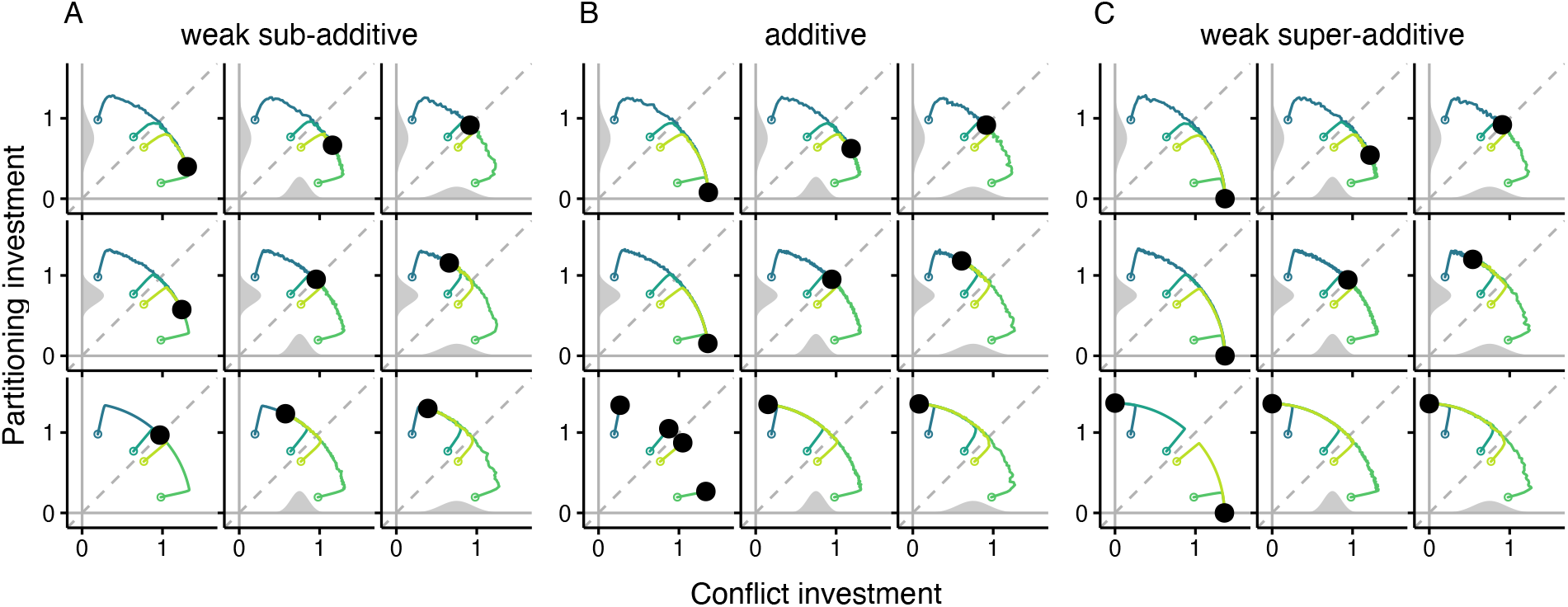
Representative simulations showing how non-heritable variation affects selection on investment for (A) weak sub-additive (*η* = −0.01), (B) additive (*η* = 0), or (C) weak super-additive (*η* = 0.01) investment costs. Each sub-panel shows the trajectories (lines) in investment space for four species started under each scenario. Open points indicate starting investments, and closed black points indicate ending investments. Gray shaded curves inside each plot show the probability density for the normal distributions used to generate non-heritable variation in conflict (bottom curve) or partitioning (left curve) trait investment evolution. No curve indicates no non-heritable variation.

### Coevolution when communities are invaded

Non-additivity of costs (*η*) and non-heritable variation (*σ_p_* and *σ_x_*) affect the possible states to which species can evolve in 2-species communities (figs. 2 and 3). To address more broadly how competitive trait coevolution affects the structure of communities, we consider the invasion of 2-species communities by a competitor that invests in either partitioning or conflict traits, and analyze the ecological (abundance) and evolutionary (trait investment) responses of the resident species and the invading species. Our focus is on the final configuration of the community, specifically the investment of species in partitioning and conflict traits, and whether this configuration is stable to the addition of new species exhibiting different investments.

A key factor needed to understand the evolution of trait investment for species is the total strength of competition it experiences, which depends on the number, abundances, and trait investments of the other species in the community. For example, if there are many other species in the community that have high population abundance and invest heavily in conflict traits, this will create strong selection for the focal species to invest in traits to decrease interspecific competition. The strength of interspecific competition experienced by species *i* is given by 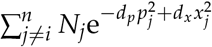. We call this the “effective competitive neighborhood”, Ω_*i*_. Analytical solutions show that the optimal investment for a species is proportional to the interspecific competition it experiences at the stationary state, denoted 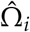 (eqs. A2–A5, fig. S4).

To investigate the effect of trait investments by both resident and invading species on community coevolution, we conducted simulations where we varied whether resident communities and the invader invest in partitioning or conflict traits in a 2×2 design. We assumed additive costs (*η* = 0) for residents and invaders. Because costs were additive, we could set whether residents invested more in partitioning or conflict traits by selecting appropriate trait starting values in the simulations. Starting with two resident species with greater investment in either partitioning or conflict traits, we first ran the simulations to stationarity for the residents. We then introduced the invader that either invested mostly in partitioning traits or in conflict traits. To allow the direct comparison between partitioning and conflict traits, we assumed that they have the same magnitude of effects on the non-focal species (*d_p_* = *d_x_* = 0.6). Because we were interested in coevolutionary patterns and not exclusion, we also chose values of *d_p_* and *d_x_* that allowed both residents and invaders to persist.

When both residents and the invader invested mostly in conflict traits (fig. 4A), the residents experienced greater effective competitive neighborhoods, which selected for an increase in their overall investment and caused a decrease in their abundance. The invader also evolved greater overall investment and had an equilibrium abundance similar to the residents. The effective competitive neighborhood for the invader changed little because the decrease in the abundance of residents offset the effect of their increased investments. When a species investing in partitioning traits invaded the same resident community (fig. 4B), investments and abundances of residents changed little. The invader, however, evolved greater investment and reached a lower abundance than the residents. The abundance and investment by the partitioning invader matched those of the conflicting invader when they invaded the same community (fig. 4A,B). When a species investing mostly in conflict traits invaded a community of partitioning residents, this increased the investment and decreased the abundance for residents, and resulted in low investment and highest abundance for the invader (fig. 4C). An invader that invested in partitioning traits invading a resident community of species investing in partitioning traits resulted in little change for all species (fig. 4D).

**Figure 4.**
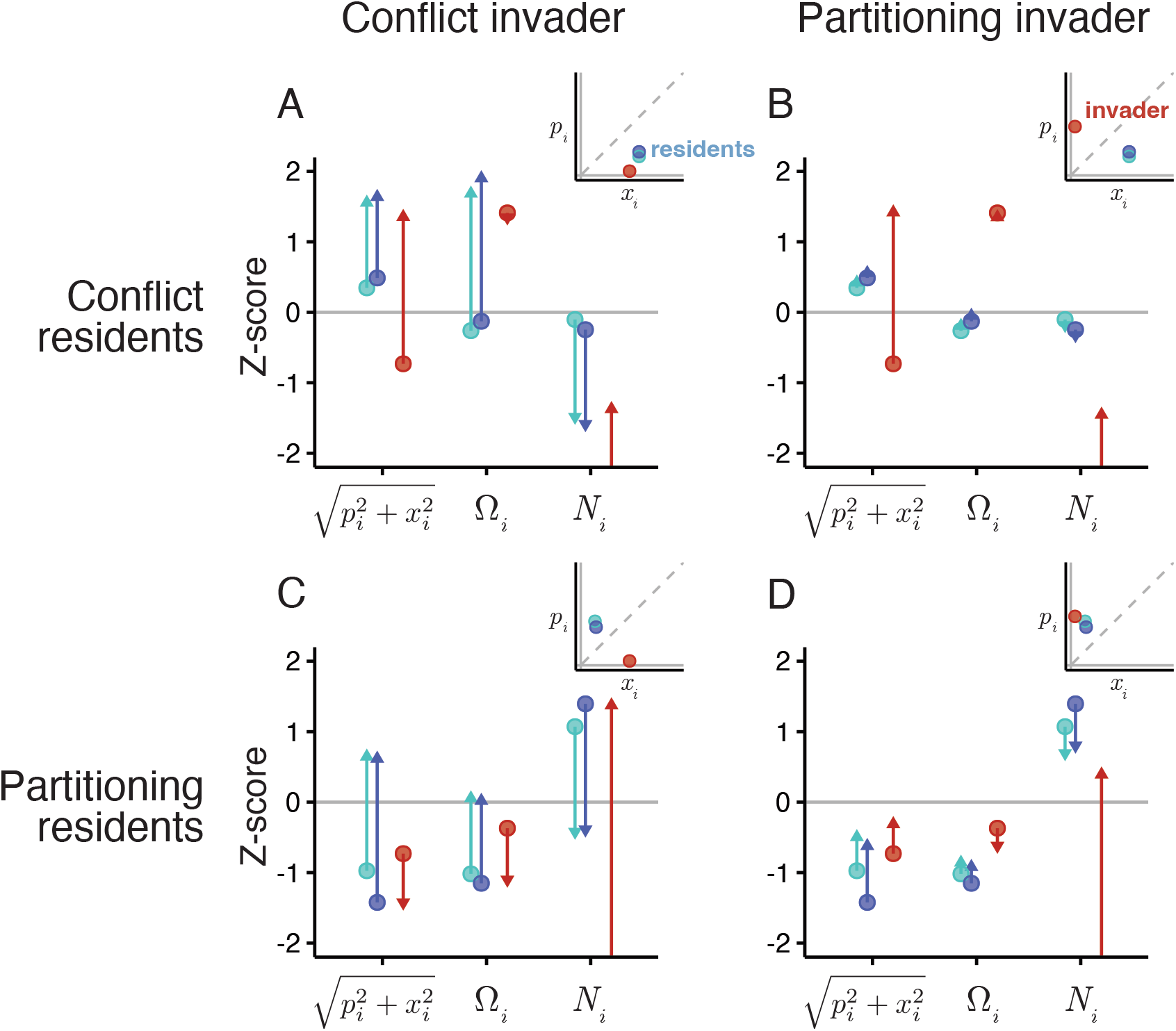
Effects of initial investments by invading and resident species on subsequent evolution of investment. Shown are z-scored total investment 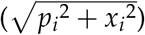, effective competitive neighborhood (Ω_*i*_), and abundance (*N_i_*) for residents (dark or light blue) of a 2-species community at stationarity that is invaded by a third species (red). Points indicate the values of each parameter when the invader was added, and arrows point to the value where the new 3-species community reached a new equilibrium. Note that invader points are the only points that do not indicate equilibrium values and thus do not conform to equations A3–A4 and equilibrium relationships described in text. Panel rows separate the relative investment of the resident species in conflict versus partitioning traits, while columns do the same for the invader. Insets show the exact investments (*p_i_* and *x_i_*) for all three species when the invader was added. Z-scoring was done within each panel (parameter set) so that they can be compared across panels. Because starting *N_i_* for the invader was always 1, we did not include this value in the z-scoring and indicated this by having the starting point outside the plot range.

Overall, these simulations show that investment in conflict traits by both invader and resident species leads to additional investment, whereas investment in partitioning traits leads to relatively less selection for changes in investment. This is true for the effects of invaders on residents (fig. 4A,C versus 4B,D) and of residents on invaders (fig. 4A,B versus 4C,D). However, these contrasting effects of investing in conflict versus partitioning traits are not necessarily symmetrical.

When species *i* invests in either type of trait, it reduces its own competition, which allows it to increase in abundance. An increase in *N_i_* increases competition for all other species in the community (i.e., it increases Ω_*j*_ for all *j* ≠ *i*). If species *i* had invested in partitioning traits, this increase in *N_i_* undercuts the competition-reducing effects of increasing *p_i_* on all other species. If species *i* had invested in conflict traits, the increase in *N_i_* exacerbates the competition-increasing effects of increasing *x_i_*. Therefore, conflict trait investment by one species increases the competition experienced by other species through its effects on both the investor’s abundance and traits, whereas when a species invests in partitioning traits, the abundance- and trait-mediated effects on other species counteract one another. This can be seen in the simulations when residents invested heavily in conflict traits and invaders in partitioning traits (fig. 4B) and when residents invested heavily in partitioning traits and invaders in conflict traits (fig. 4C), where we found that the species investing in conflict traits had a greater influence on investments than those investing in partitioning traits. Thus, in communities with species expressing both partitioning and conflict traits, we expect the evolution of conflict traits to be the stronger driver of trait evolution for all species.

### Competitive exclusion

In our model, species that invest more in conflict traits are generally more abundant and invest less overall than species investing mostly in partitioning traits (fig. 4). If an invader invested heavily in conflict traits—the type most likely to cause competitive exclusion—arrives to a community, greater total investments by residents help shield them from the new source of competition, whereas greater resident abundance should slow invader population growth and provide the resident a longer time for evolutionary rescue via greater investment. Our next simulations assess how these factors together affect the likelihood of an invader excluding resident species that invest in either partitioning or conflict traits.

We simulated the invasion of 2-species equilibrium communities as in the previous section, but with one resident species always invested more in partitioning traits (partitioning resident: *p_t_* ≫ *x_t_*) and the other always invested more in conflict traits (conflict resident: *p_t_* ≪ *x_t_*). We varied the strength of effects of trait investment on other species (*dp* and *dx*) because these parameters shape interactions between residents and invaders. We varied the strength of the effect of investing in traits on other species (*d_p_* and *d_x_*) because these parameters shape interactions between residents and invaders, but we focused on *d_p_* since this had the strongest effect on the coevolutionary dynamics (fig. S5). At equilibrium, the conflict resident invested less overall and was more abundant than the partitioning resident, and these disparities increased with *d_p_* (fig. 5A,B). Specifically, when *d_p_* = 0.3, the conflict resident moderately invested, while it invested very minimally when *d_p_* = 0.7; the portioning resident maintained high investment in both scenarios (fig. 5C,D). When an invader investing in conflict traits (conflict invader: *p*_*t*=0_ = 0, *x*_*t*=0_ = 1) was added to the equilibrium 2-species community, overall investment was more important than abundance in determining which resident was excluded. When the conflict resident invested moderately (*d_p_* = 0.3; fig. 5C) at the 2-species equilibrium, the addition of the invader caused only a temporary drop in the conflict resident’s abundance in response to the increase in the effective competitive neighborhood (fig. 5E). In contrast, when the conflict resident invested minimally (*d_p_* = 0.7; fig. 5D), it went extinct as it was unable to increase its investment rapidly enough in response to the increase in the effective competitive neighborhood (fig. 5F). Because the partitioning resident always invested more overall, the effects of the conflict invader were less severe and did not cause it to go extinct. This pattern where the conflict resident was easier to exclude than the partitioning resident was constant over a wide range of *d_p_*, *d_x_*, and the conflict invader’s starting total investment (fig. S6).

**Figure 5.**
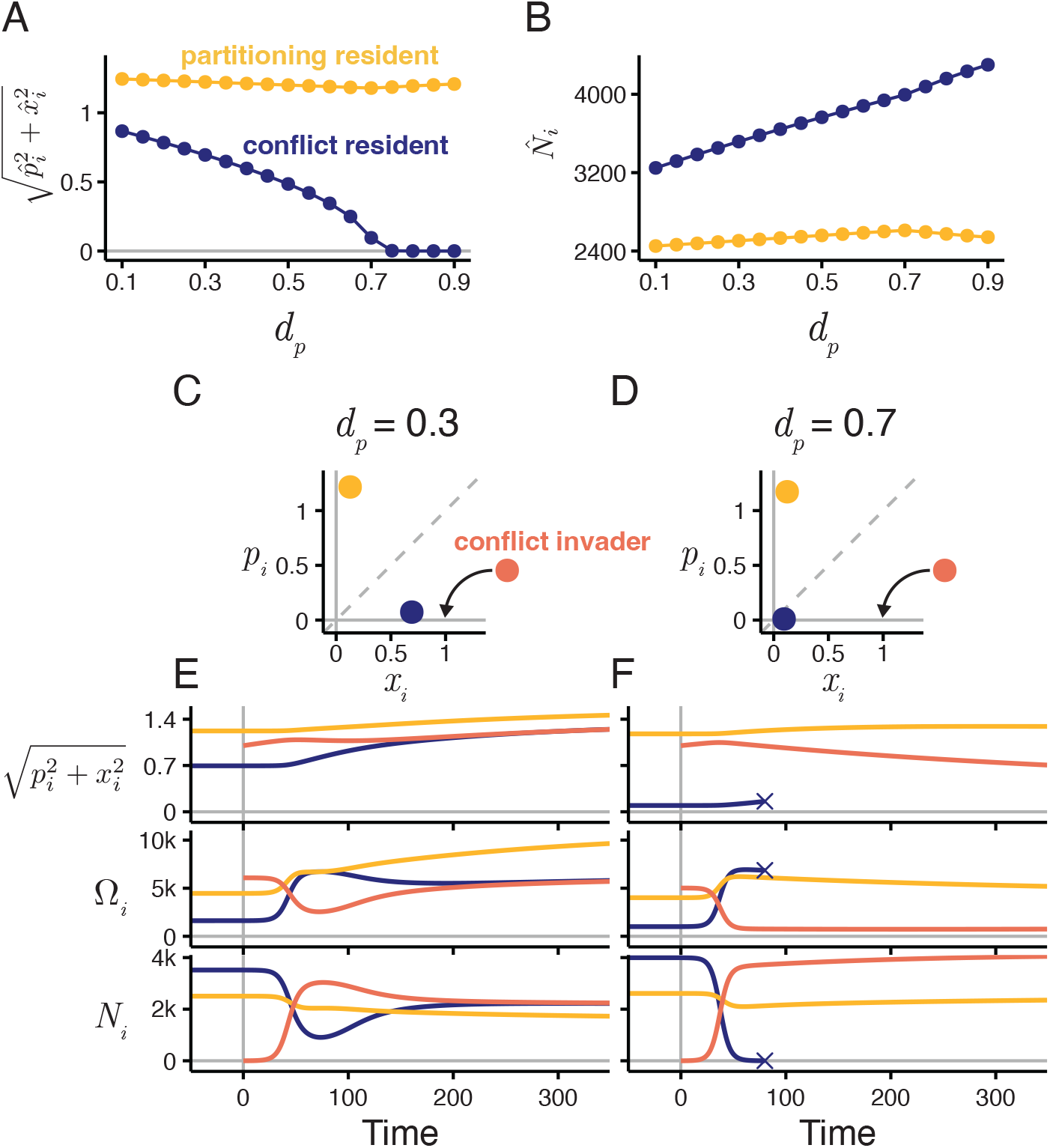
Factors affecting exclusion of residents in a 2-species mixed community. For the equilibrium 2-species resident community, (A) total investment 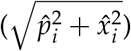 and (B) abundance 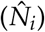 by the species investing more in partitioning traits (“partitioning resident”; gold) and by the species investing more in conflict traits (“conflict resident”; purple) depend on how strongly partitioning investment affects the competition experienced by other species (*d_p_*). (C) Equilibrium investments for the 2-species community when *d_p_* = 0.3 and (D) when *d_p_* = 0.7, with investments of the invader species shown in orange. (E,F) Total investment 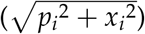, effective competitive neighborhood (Ω_*i*_), and abundance (*N_i_*) for both residents and an invader invested in conflict traits (“conflict invader”; orange) added to the community from (E) panel C or (F) panel D. Crosses indicate extinction.

These simulations show that investing in conflict traits can make species more likely to be excluded from a community that also contains species that invest in partitioning traits. By reducing other competitors’ abundances, thereby selecting for reduced investment for itself and increased investment for others, competitors that invest proportionally more in conflict traits make themselves the more vulnerable to exclusion despite being the more abundant in the community. We might therefore expect higher turnover and greater variation in abundance for species invested in conflict traits. This bias in exclusion risk could result in communities that, on average, contain fewer species investing strongly in conflict traits.

## Discussion

Here, we showed how coevolution shapes the evolution of traits that govern the strength of competition between species. Two types of competition traits have distinct effects on the rate and outcome of coevolution. Conflict traits increase the impacts of competition on other species while simultaneously reducing competition for the evolving species. In contrast, partitioning traits simultaneously reduce the strength of competition experienced by the species evolving the traits and the species with which it interacts. The level of investment in conflict and partitioning traits depends on whether costs for investing simultaneously in both types of traits are additive, sub-additive, or super-additive. If the trade-off between partitioning and conflict traits is sub-additive or additive, we would expect species to invest in both, whereas super-additive costs should result in investment strategies of only partitioning or only conflict traits. Furthermore, non-heritable variation acts to make the effective trade-off more sub-additive; additive and even weakly super-additive trade-offs become effectively sub-additive when there is sufficient non-heritable variation, resulting in simultaneous investment in both trait types. Thus, the nature of investment costs, and how they are modified by non-heritable variation, determine whether species evolve a mixed strategy comprising both partitioning and conflict traits (sub-additive) or evolve a specialist strategy of only partitioning or only conflict traits (super-additive). In both cases, however, we would expect that communities should always contain mixed investments, either by all species showing mixed strategies or by different species specializing on different strategies. Although partitioning and conflict traits should occur simultaneously in communities, they affect community coevolution in different ways. The evolution of conflict traits by a species causes both increases in the abundance of the evolving species (by reducing competition it experiences) and increases in the per capita strength of competition it exerts on other species. Therefore, the evolution of conflict traits leads to greater ecological (abundance) and evolutionary (traits) impacts on a community than partitioning traits.

Despite the ubiquity of conflict traits, their impacts on competitive communities are relatively understudied, partly because the question of how so many species can coexist in nature is assumed to be answered by resource partitioning. However, some empirical evidence has shown that exclusion-promoting traits, not necessarily niche partitioning, often evolve in response to changes in competition (Germain et al., 2020; Hart et al., 2019; Miller et al., 2014;Zhao et al., 2016). Furthermore, recent work on annual plant communities found that average fitness shows greater phylogenetic signal than niche partitioning (Godoy et al., 2014) and that many commonly measured functional traits correspond more closely to competitive dominance than to niche differences (Kraft et al., 2015). These together suggest that conflict traits that reduce competition for a focal species at the expense of its competitors might be important for the evolution of competitive communities. Our results support this hypothesis and show that trait evolution for entire communities should be driven most strongly by investments in conflict traits.

Although investors in conflict traits should drive greater investments in other species, their suppression of competitors’ abundances combined with a cost to investment causes these conflict species themselves to evolve reduced investment. Conflict-investing species invading a community should therefore experience evolutionary disarmament after the initial period of ecological suppression of competitors. On the surface, this appears to contradict two major hypotheses related to invasive species evolution. First, the novel weapons hypothesis predicts the evolution of stronger armament after the invasion because of selection to suppress native competitors (Callaway and Ridenour, 2004; Inderjit et al., 2006). Second, the evolution of increased competitive ability (EICA) hypothesis predicts evolution of greater competitive dominance of invaders because they are released from selection due to natural enemies (Blossey and Notzold, 1995). However, our simulations do show small evolutionary increases in investments by invaders upon arrival (fig. 5E,F), but this is followed by a slower and stronger decrease after native species are suppressed. This suggests a secondary stage of invasive species competitive coevolution, where they evolve reduced suppressive traits when natives are rare enough that the costs of maintaining those traits are no longer outweighed by the benefits.

These two stages of invader evolution, first to increase investment in competitive traits and then to decrease investment, may also play out through space as the invasive species coevolves with natives that are more abundant at the edge of the invader’s range (Miller et al., 2020;Thompson, 2005). Empirical evidence for this comes from garlic mustard (*Alliaria petiolata*), a widespread Eurasian invader of North American forest understories. Garlic mustard produces allelopathic compounds that suppress heterospecific plants but are costly when only competing with conspecifics (Evans et al., 2016). A number of studies have shown that its allelopathic effects on North American competitors decrease with time after invasion and that this pattern is consistent with evolution via selection for reduced allelopathy (Bossdorf et al., 2004; Evans et al., 2016; Huang et al., 2018; Lankau et al., 2009). For costly traits that effectively suppress native species, this secondary stage of disarmament might be common.

Another effect of evolutionary disarmament by conflict-trait investors is that these species should have the greatest likelihood of being excluded from a community when it is invaded by another species showing conflict traits. This “what goes around comes around” phenomenon makes communities containing many species with conflict traits eco-evolutionarily transitory. It also means that native species that suppress others but have gone through disarmament might be most vulnerable to exclusion, despite being more abundant than their competitors. Like many eco–evolutionary processes (Losos and Ricklefs, 2009), the impermanence of heavy conflict investors might be most obvious on remote islands. With long periods of evolutionary disarmament in the absence of new colonizations, repeat invasions by conflict investors from the competitor-rich mainland to competitor-poor islands could produce cycles of invasion, dominance, disarmament, and extinction. This is similar to “taxon cycles,” where species go through repeated periods of greater dispersal to islands, followed by evolved specialization and range fragmentation once they establish on islands that ultimately cause them to go extinct (Losos and Ricklefs, 2009; Wilson, 1961). The evidence for taxon cycles is mixed (Losos, 1990, 1992; Mayr and Diamond, 2001; Taper and Case, 1992) but strongest for birds in the West Indies (Ricklefs and Bermingham, 2002) and for Melanesian ants (Darwell et al., 2020; Economo and Sarnat, 2012; Wilson, 1961). One of the main observations made to support taxon cycles is the transition of recent invaders being abundant and widespread to older taxa having a greater risk of extinction (Ricklefs, 2010). This is also consistent with serially invading conflict investors followed by disarmament and may indicate that this, along with other processes such as coevolution of natural enemies (Ricklefs and Bermingham, 2002), might underlie some of the episodic extinction–colonization events that exemplify taxon cycles.

## Conclusions

In trying to understand how trait evolution affects species coexistence, research most often focuses only on traits promoting coexistence, such as those generating resource partitioning. Here, we show that communities are likely to contain evolutionary investments in both coexistence- and exclusion-promoting traits, and that the latter should have the greatest community-wide effects on abundances and investments in competitive traits. Because traits promoting exclusion suppress the abundance of competitors, in the long run selection should favor reduced investment in species possessing these traits. In turn, lower investment corresponds to a higher risk of later exclusion by an invading species. Whether coevolution promotes coexistence or exclusion may depend on the competition traits possessed by a given species. Coevolution may provide an open door for those that play nice and a revolving door of exclusion for those that do not.

## Appendix A: Analytical solutions, non-negative investments, and non-heritable variation

### Analytical solutions

Below are the full equations for fitness and investments at time *t* + 1, with the parameter for effective competitive neighborhood (Ω_*i*_) defined as the total community abundance accounting for the effect of species’ investments on competition experienced by species *i*. These equations are used for the analytical solutions in this section, and in these solutions, equilibrium parameters are indicated by circumflexes (e.g., 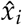 for *x_i_*).

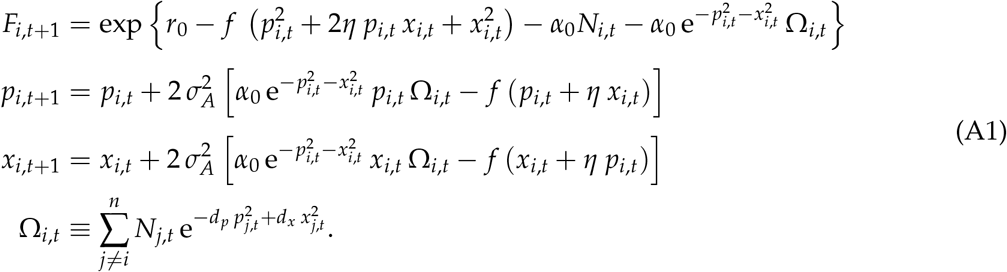

For a 1-species community at equilibrium, unless *η* = −1,

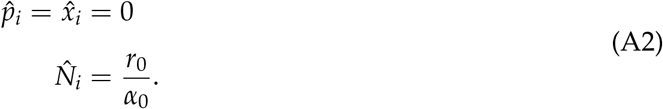

Because investments cannot be negative and

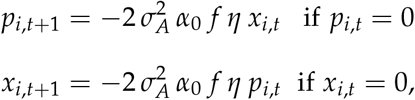

*p*_*i,t*+1_ and *x*_*i,t*+1_ will remain at zero unless *η* ≥ 0 or the other investment type is zero. This means that equilibria where exactly one investment is at zero only coincide with additive or super-additive costs. For these two equilibria,

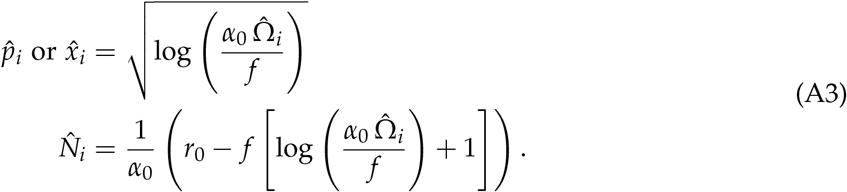

These are the only possible equilibria for super-additive costs. However, if costs are additive, there are infinite equilibria along a neutrally stable ring defined by the overall distance from the origin:

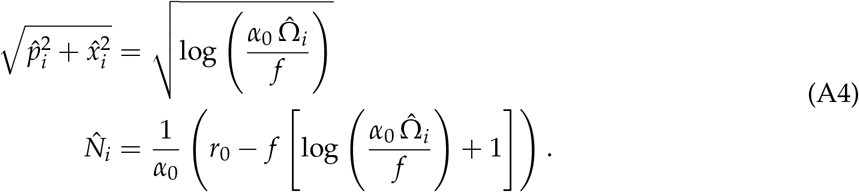

For sub-additive costs, we have one equilibrium where 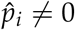 & 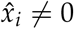:

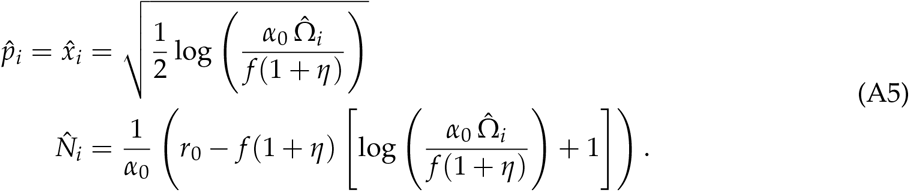

The equilibria shown in equations A3–A5 are undefined for low values of 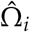, in which case species evolve to 0 investment. This will occur when

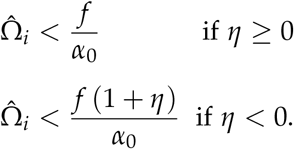

### Keeping investments non-negative

To keep investments ≥ 0, we used the ramp function *R*(*z*) = max(0, *z*). We used a ramp function instead of absolute values because the latter causes fluctuations in the investments when they approach zero (they “bounce off” the zero bound) that persist for extended periods. A more important disadvantage is that derivatives of these functions are undefined at zero. The derivative of the ramp function is the Heaviside step function:

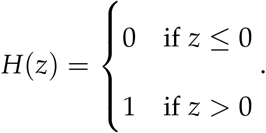

Thus, all investment derivatives of the form *∂x/∂z* and *∂p/∂z* are changed to *H*(*p*(*z*)) *∂p/∂z* and *H*(*x*(*z*)) *∂x/z*, respectively, to account for investments being non-negative.

### Effects of non-heritable variation on investment evolution

When there is non-heritable variation in investment evolution, the second order Taylor series approximation for the expected value of *x*_*t*+1_ is

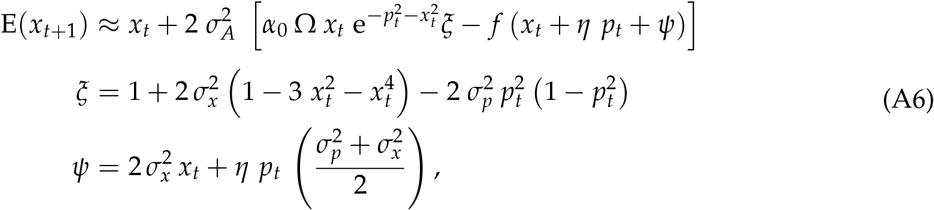

where we drop the index for species to reduce clutter.

## Appendix B: General matrix forms of equations and Jacobian

### Matrix forms of equations

This section defines a general form of the equations in the main text, for any *q* suites of traits, using matrix notation. We used this form for the derivatives in the subsequent section, and we believe that keeping them in a more general form will help make them more useful to those interested in building from this work. Here, ^T^ indicates transposition, multiplication between matrices is always matrix multiplication, bold capital letters indicate a matrix, and bold lowercase letters indicate a vector. See Table B1 for descriptions of the parameters.

**Table B1.** Parameters used in the matrix equations.

| Parameter | Description |
| --- | --- |
| $q$ | The number of trait suites in which species can invest. |
| $n$ | The number of species in the community. |
| $\mathbf{v}_i$ | A $q \times 1$ vector containing the investments for species $i$. |
| $F_i$ | Per-capita growth for species $i$. |
| $N_i$ | Population density of species $i$. |
| $\alpha_0$ | Base density dependence. |
| $r_0$ | Baseline growth rate. |
| $f$ | Cost of increasing investments on the growth rate. |
| **C** | Contains parameters controlling non-additive costs. All values on the diagonal are 1. |
| $\eta_{k,l}$ | Non-additive cost of increasing both the $k^{\text{th}}$ and $l^{\text{th}}$ suites of traits. When $\eta_{k,l} > 0$, investing in both suites incurs an extra cost compared to investing in each separately. When $\eta_{k,l} < 0$, investing in both suites is less costly. Non-additive trade-offs are symmetrical (i.e., $\eta_{k,l} = \eta_{l,k}$ for all $l$ and $k$). |
| **D** | Contains parameters that determine how investment in one species affects others. All off-diagonal elements are zero. |
| $d_k$ | Effect on other species when one invests in the $k^{\text{th}}$ suite of traits. When $d_k < 0$, this investment increases competition experienced by all other species (i.e., the $k^{\text{th}}$ traits are conflicting). Alternatively, when $d_k > 0$, the same investment decreases the effect of competition on all species (i.e., the $k^{\text{th}}$ traits are partitioning). |
| $\sigma_A^2$ | Additive genetic variance. |

The following 4 equations correspond to Equations 1–4 from the main text.

The per-capita growth for species *i* is

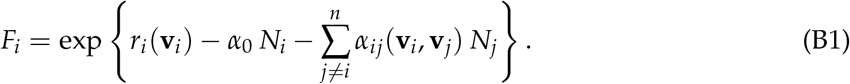

The parameter *r_i_*(**v***_i_*) describes the direct costs of investing to the growth rate:

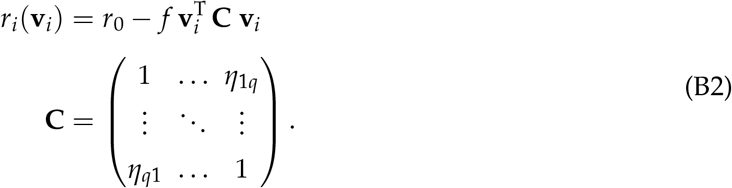

The term *α_ij_*(**v***_i_*, **v**_*j*_) in equation B1 represents how investments influence the effects of interspecific competition:

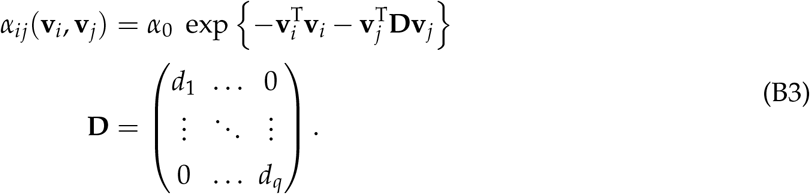

Investments at time *t* + 1 are

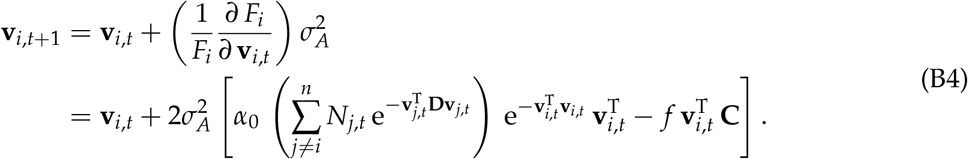

### Jacobian

The Jacobian is

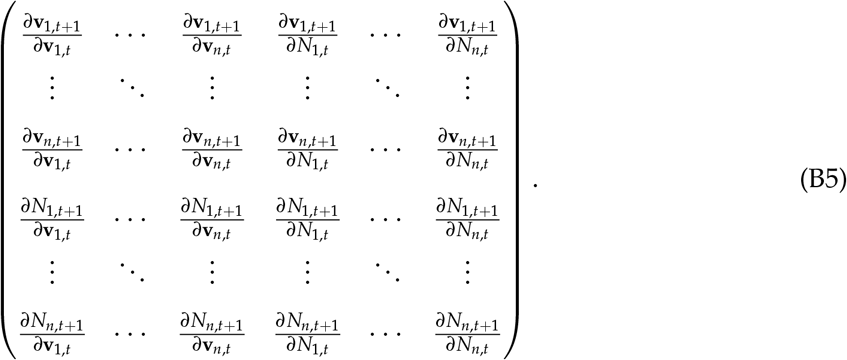

It is made up of the following derivatives:

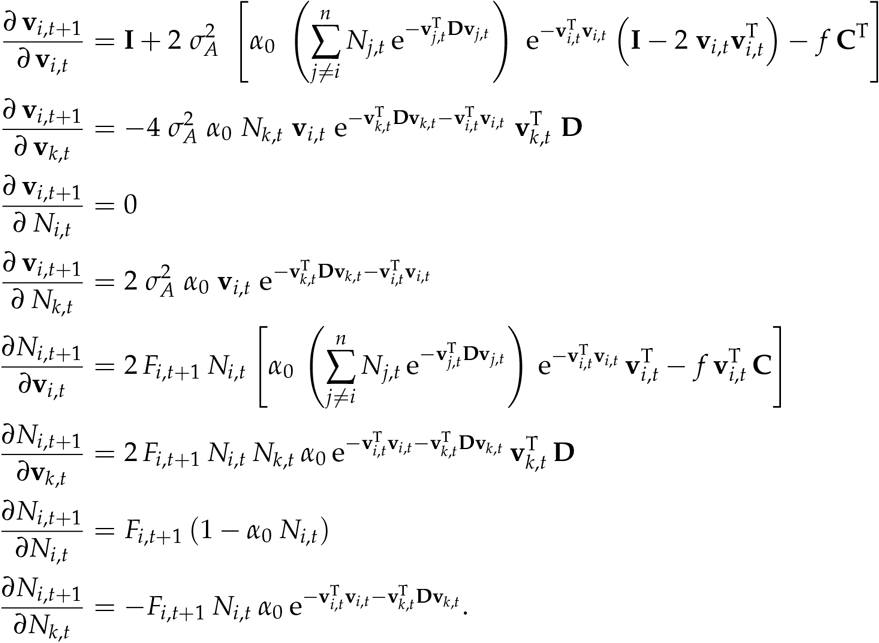

