## Supplemental Material for "When should coevolution among competitors promote coexistence versus exclusion?"

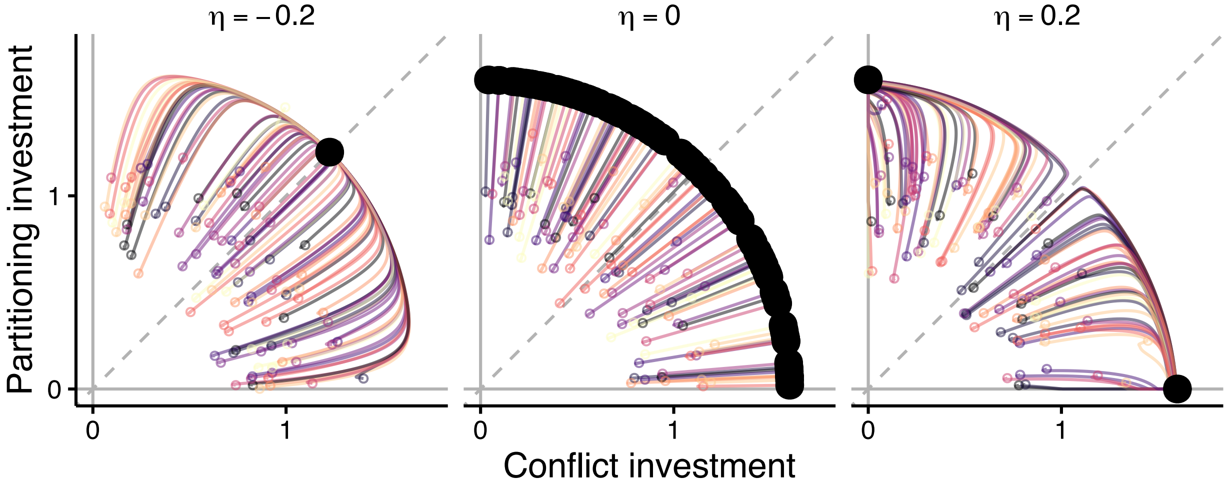


**Figure S1.** Evolutionary trajectories for simulations of 10-species communities with varying $\eta$. Shown are 10 simulations (indicated by color) where species' starting investments were generated randomly near a ring defined by $\sqrt{p_{i,t=0}^2 + x_{i,t=0}^2} = 1$. The total investment was drawn from a normal distribution (truncated above zero) with mean of 1 and standard deviation of 0.2, while the angle along the origin was generated from a uniform distribution from 0° to 90°. Small open points indicate starting investments, and large closed points indicate the points to which species evolved. To highlight just the effects of $\eta$, both sets of traits were nearly neutral ($d_p = d_x = 10^{-6}$), as for Figure 2.

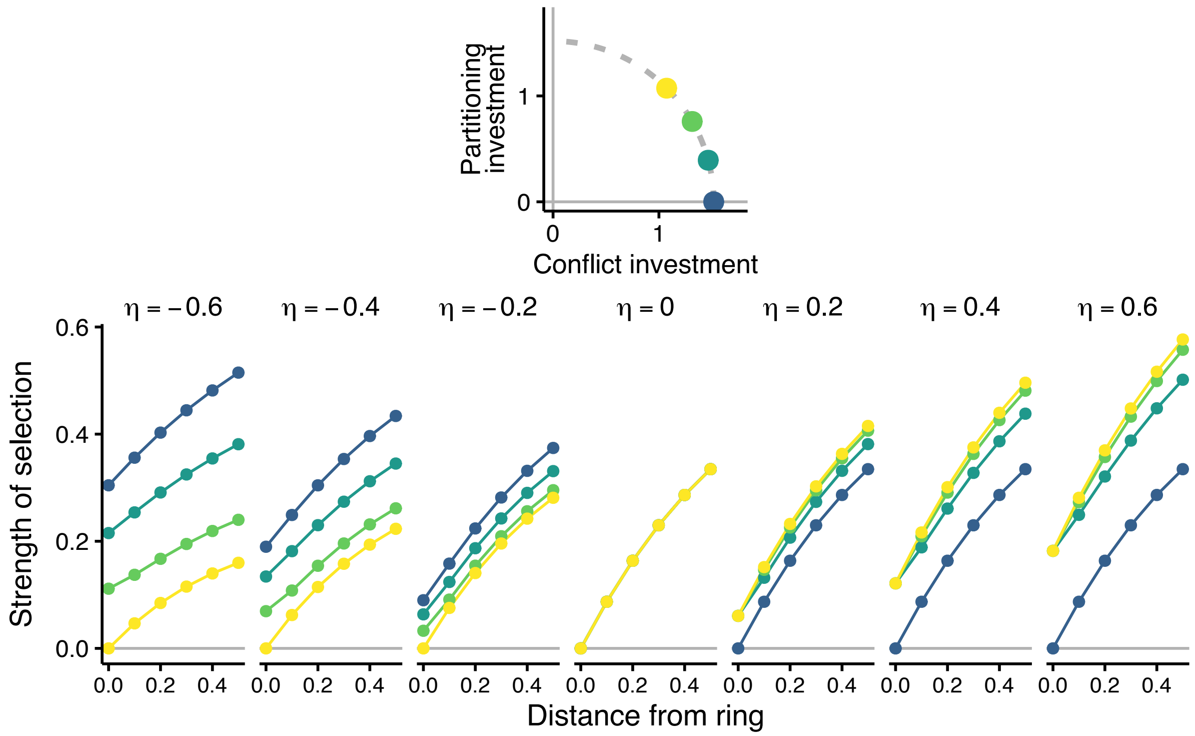


**Figure S2.** Strength of selection ($\sqrt{(F_i^{-1}\,\partial F_i\,/\,\partial p_i)^2 + (F_i^{-1}\,\partial F_i\,/\,\partial x_i)^2}$) in relation to distance from a ring representing the total investment at equilibrium ($\sqrt{\hat{p}_i^2 + \hat{x}_i^2}$) and the location along this ring. The ring radius was calculated using equations A3–A5 with an equilibrium effective competitive neighborhood ($\hat{\Omega}_i$) of 10,000. The point colors in the main plot correspond to the positions along the ring in the upper inset. The yellow point is the equilibrium location when $\eta < 0$, the darkest green point is equilibrium when $\eta > 0$, and any of the points are equilibria when $\eta = 0$. Panel columns separate the degree of investment cost non-additivity ($\eta$).

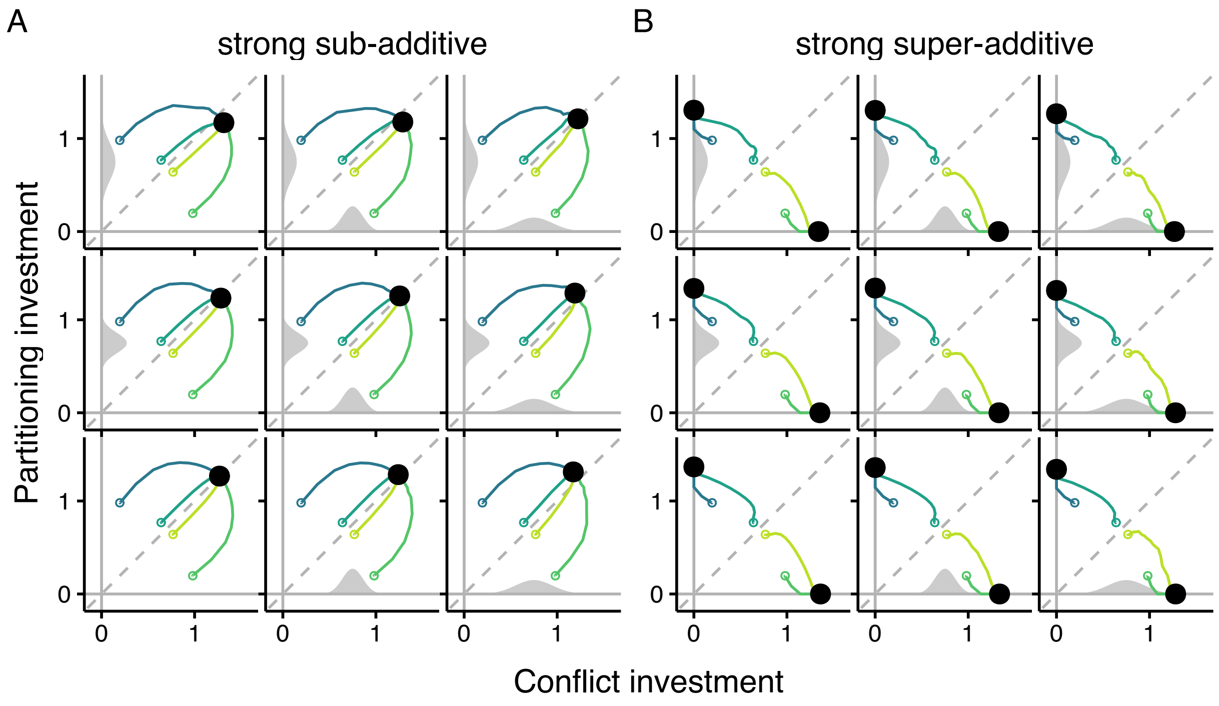


**Figure S3.** Representative simulations showing that weakening selection does not significantly affect investment evolution when investment costs are (A) strongly sub-additive ($\eta = -0.6$) or (B) strongly super-additive ($\eta = 0.6$). Each sub-panel shows the trajectories (lines) in investment space for four species started under each scenario. Open points indicate starting investments. Closed, black points indicate ending investments. Gray shaded curves inside each plot show the probability density for the normal distributions used to generate non-heritable variation in conflict (bottom curve) or partitioning (left curve) trait investment evolution. No curve indicates no non-heritable variation.

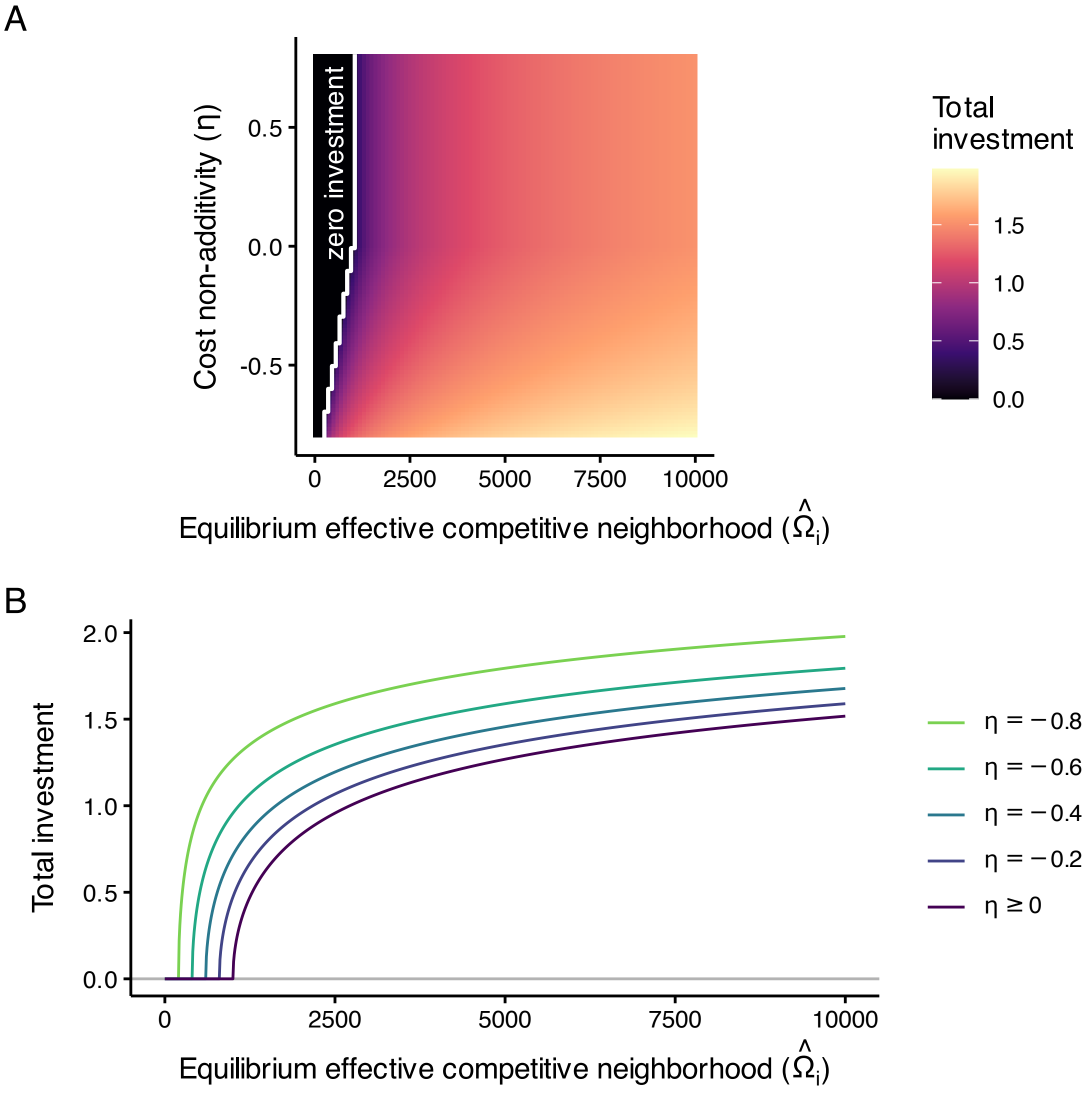


**Figure S4.** Effects of cost non-additivity ($\eta$) and effective competitive neighborhood at equilibrium ($\hat{\Omega}_i$) on total investment at equilibrium ($\sqrt{\hat{p}_i^2 + \hat{x}_i^2}$) by focal species i, based on Equations A3–A6. (A) Heatmap showing total investment (color) by $\eta$ and $\hat{\Omega}_i$. The white line separates the areas with zero and non-zero total investment. (B) Effect of $\hat{\Omega}_i$ on total investment, where line color indicates $\eta$. Because equilibrium investments when $\eta \geq 0$ do not depend on $\eta$ (Equations A3, A4), the relationship between $\hat{\Omega}_i$ and total investment is the same for all $\eta \geq 0$.

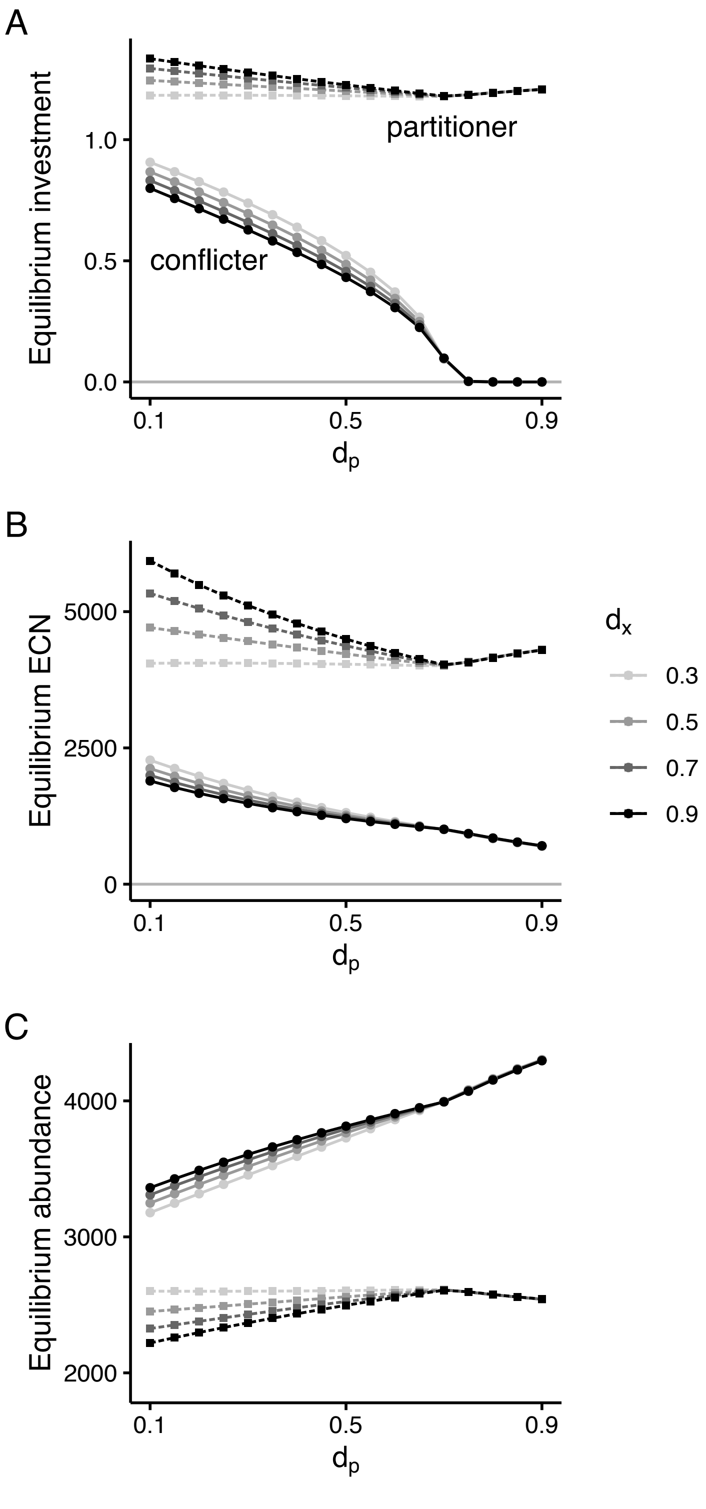


**Figure S5.** Equilibrium total investment ($\sqrt{\hat{p}_i^2 + \hat{x}_i^2}$) effective competitive neighborhood ("ECN," $\hat{\Omega}$), and abundance ($\hat{N}$) for 2-species communities where one species invests heavily in partitioning traits ("partitioner," squares and dotted lines) and another in conflict traits ("conflicter," circles and solid lines). The x-axis shows how strongly partitioning investment affects the competition experienced by other species ($d_p$). Point and line color indicate the effect of conflict investment on other species ($d_x$).

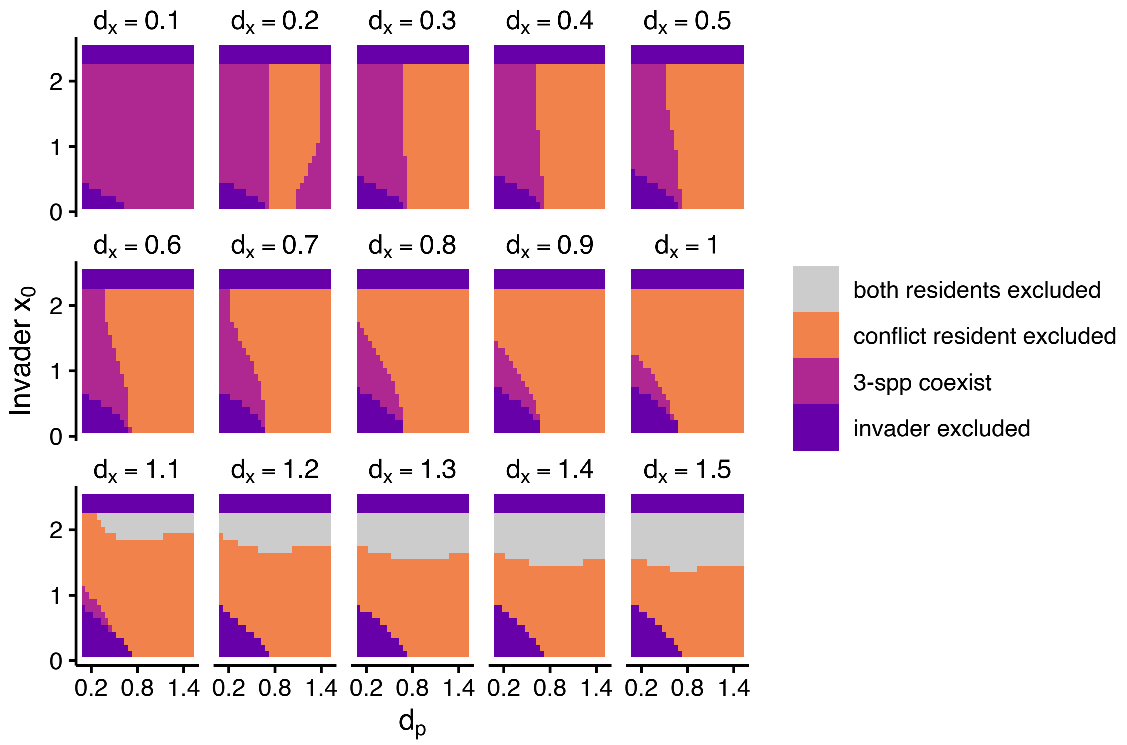


**Figure S6.** Outcomes from adding a conflict invader to a 2-species community where one resident invests heavily in conflict traits and the other in partitioning traits. Each pixel in the panels represents the outcome from a single simulation. Four outcomes were possible: the invader excluded both residents (gray), the conflict resident was excluded (orange), all 3 species coexisted (magenta), or the invader was excluded (purple); the partitioning resident was never the only species excluded. The x-axis is the effect of partitioning investment on competition experience by other species ($d_p$), and the y-axis is the starting conflict investment by the invader; in all simulations, the invader did not invest in partitioning traits. Panels separate the effect of investment in conflict traits on other species ($d_x$). Equilibrium communities varied by $d_p$ and $d_x$, so each column within a panel indicates invasions of the same community by different invaders. Each row (across all panels) indicates outcomes caused by the same invader to different equilibrium communities. Three-species coexistence with high $d_p$ and $d_x = 0.2$ is not highlighted in the main text because it represents an edge case where the conflict resident is nearly extinct ($N_i < 10$) but maintained by the strong investment by the partitioning species for 4,000 time steps while it slowly evolves greater investment. This behavior disappears with small changes in $d_x$.
